# Inhibition of Myoglobin Oxidation by Food Grade Botanical Antioxidant Formulations

**DOI:** 10.1101/2024.11.11.622983

**Authors:** Valentina Vergara-Stange, Ana Batista-Gonzalez, Rodrigo A. Contreras

## Abstract

This study explores the antioxidant capabilities of green tea and acerola extracts in inhibiting myoglobin oxidation, a critical factor in preserving color stability in food products, mainly plant-based meat analogs. Extracts were analyzed for their antioxidant activity using DPPH and ORAC assays, followed by evaluations of their inhibitory effects on myoglobin oxidation at 25 °C and 35 °C. Acerola showed quick inhibitory effects even at lower concentrations and moderate temperatures, likely due to its high ascorbic acid levels. On the other hand, green tea’s effectiveness varied with temperature, with inhibition rates rising at 35 °C, probably because its polyphenolic compounds are sensitive to heat. The kinetics and thermodynamics analysis showed positive activation energy (Ea) for green tea, meaning it becomes more active as temperatures rise. In contrast, acerola’s negative Ea indicates heat reduces its efficacy. These results suggest that acerola and green tea could be natural alternatives to synthetic antioxidants, helping preserve the appearance and stability of different food products.

## 1. Introduction

Antioxidants are vital bioactive compounds that help prevent oxidation in food and pharmaceutical products, extending their shelf life while preserving quality and safety. In the food industry, antioxidants are crucial in slowing down the spoilage and rancidity of fats and oils. By preserving the color, flavor, and nutritional value of food products, antioxidants help extend the shelf life of processed foods and reduce waste (Girish et al., 2023). Today, one of the main challenges the food industry faces is the rapid spoilage of products, leading to significant financial losses and environmental issues due to increased food waste (Leyva-Porras et al., 2021).

Food spoilage occurs through several mechanisms, primarily microbial growth, oxidation, and enzymatic self-decomposition (Leyva-Porras et al., 2021). Among these, oxidation is particularly problematic as it can be triggered by factors such as UV light, temperature changes, pH shifts, and oxygen exposure (Bensid et al., 2022). This process generates reactive oxygen species (ROS), including hydroxyl radicals (•OH), superoxide anions (O□•□), hydroperoxyl radicals (•OOH), and hydrogen peroxide (H□O□), which lead to further oxidation and food degradation (Poljsak et al., 2021).

Synthetic and natural antioxidants are commonly used in the food industry to combat oxidation. However, synthetic antioxidants like butylated hydroxytoluene (BHT), butylated hydroxyanisole (BHA), and tertiary butylhydroquinone (TBHQ) have raised health and safety concerns, leading regulatory authorities to limit their use (Xu et al., 2021). This has pushed consumers to seek natural alternatives, as they perceive them to be safer and more sustainable (Gutiérrez-del-Río et al., 2021)

Beyond preservation, natural antioxidants from fruits, vegetables, and herbs have become increasingly popular due to their health benefits. They often contain beneficial bioactive compounds such as polyphenols, carotenoids, vitamins, and minerals, contributing to overall well-being (Knez et al., 2024).

Natural antioxidants have additional advantages. They are generally abundant, affordable, and considered safe for consumption. These qualities make natural antioxidants a sustainable alternative for the food industry, aligning with the current consumer trend toward eco-friendly and health-conscious products (Ghosh et al., 2022; Pappalardo et al., 2020). Alongside the preference for natural ingredients, there is a growing shift towards plant-based diets driven by health benefits and a lower environmental footprint (Carey et al., 2023).

However, the plant-based food industry faces unique challenges. It is not just about replacing animal proteins; to be widely accepted, plant-based products must also mimic the sensory qualities of meat, such as texture, flavor, color, and aroma (Jang & Lee, 2024). To appeal to traditional meat consumers, plant-based meat alternatives must closely replicate the look and taste of real meat before and after cooking (Ahmad et al., 2022). This has led to the use of innovative ingredients like non-animal myoglobin and leghemoglobin, which help recreate the appearance of meat (Wilson et al., 2024).

However, using myoglobin in food formulations introduces new challenges. Myoglobin, a protein that binds oxygen, gives muscle tissue its red color, which consumers find visually appealing. But when myoglobin loses its bound oxygen, its heme iron becomes exposed, making it prone to oxidation. Consumers often associate a brownish color with spoilage or lower quality (Grunert et al., 2004). Once myoglobin oxidizes, food products containing it lose their visual appeal. Therefore, stabilizing myoglobin to prevent oxidation is crucial for traditional and plant-based food products, as it ensures the product remains visually attractive and has a longer shelf life (Muñoz-González et al., 2022).

One effective strategy to stabilize myoglobin is the addition of antioxidants. Both synthetic and natural antioxidants have been explored for this purpose. Among natural options, extracts from fruits and vegetables rich in polyphenols, carotenoids, tocopherols, and ascorbic acid have shown promise in preventing myoglobin oxidation (Su et al., 2024).

As the demand for plant-based and natural ingredients grows, the food industry is increasingly interested in botanical sources of antioxidants. These include extracts from various fruits, vegetables, herbs, and spices, offering a broad range of compounds that fight oxidation. Polyphenolic compounds such as flavonoids, phenolic acids, and tannins are potent due to their high antioxidant capacity. Additionally, carotenoids and vitamins C and E help neutralize ROS, providing oxidative stability that preserves food product quality and shelf life (Petcu et al., 2023). As a result, plant-derived antioxidants are seen as a promising, natural solution for oxidative stability in food products.

This study addresses the growing need for effective antioxidant formulations that stabilize myoglobin in food applications. Specifically, it focuses on food-grade botanical extracts rich in polyphenols, carotenoids, anthocyanins, betalains, and vitamins—compounds that have shown strong antioxidant activity in previous studies. By evaluating a range of botanical extracts and their effectiveness at preventing myoglobin oxidation under various temperature conditions, this research seeks to provide insights into natural solutions for maintaining color and quality in food products.

The study tests the antioxidant activity of nine freeze-dried fruit and vegetable extracts or juice powders using various *in vitro* assays. These extracts will then be evaluated for their ability to inhibit oxidation in equine myoglobin—a model system commonly used in food science due to its structural and functional similarity to animal myoglobin. Testing under different temperature conditions will help determine how these antioxidant formulations perform under typical storage and processing scenarios.

In summary, this study aims to contribute to developing natural antioxidant solutions for stabilizing myoglobin in both plant-based and traditional food products. By focusing on botanical sources, the research aligns with consumer preferences for natural, sustainable ingredients and addresses the technical challenges of maintaining color stability in food products. The findings are expected to support creating visually appealing products with extended shelf life, benefiting both the food industry and environmentally conscious consumers.

## 2. Materials and methods

### 2.1. Reagents and chemicals

2,2-diphenyl-2-picrylhydrazyl (DPPH•) and 6-hydroxy-2,5,7,8-tetramethylchroman-2-carboxylic acid (Trolox®) were purchased from Calbiochem (San Diego, California, USA), 2,2’-azobis(2-methylpropionamidine) dihydrochloride (AAPH), pyrogallol red (PGR) and equine heart myoglobin were purchased from Sigma-Aldrich (St. Louis, MO, USA). Ferrozine was purchased from Merck (Darmstadt, Germany). Iron (II) sulfate (FeSO_4_) and bovine serum albumin (BSA) were purchased from Winkler (Santiago, RM, Chile). Ethylenediaminetetraacetic acid (EDTA) was purchased from VWR (Radnor, PA, USA).

### 2.2. Food-grade botanicals

The botanicals used in this study were sourced from local suppliers. They included green tea (*Camellia sinensis*) extract, açaí (*Euterpe olearacea*) juice powder, acerola (*Malpighia emarginata*) juice powder, beetroot (*Beta vulgaris*) juice powder, rosemary (*Salvia rosmarinus*) extract, and a blend of rosemary and green tea extract. All ingredients were dissolved in double-distilled water except for the green tea, which was dissolved in 30% ethanol. The final concentrations ranged from 0 to 10 mg/mL, depending on the assay.

### 2.3. LC-MS/MS Phytochemical Analysis of Botanicals

The botanical extracts, green tea, acerola, rosemary, and a rosemary-green tea blend, were analyzed in triplicate using a UHPLC system coupled to a Bruker compact QTOF mass spectrometer at the Omics Technologies Unit’s Mass Spectrometry Service, Faculty of Biological Sciences, Pontificia Universidad Católica de Chile. Chromatographic separation was performed on a Kinetex column (2.1 mm × 100 mm, 1.7 µm, Phenomenex), with a mobile phase consisting of 0.1% formic acid in water (solvent A) and 90% acetonitrile with 0.1% formic acid (solvent B). The gradient was set as follows: 0.0–0.5 min, 88% A; 0.5– 11.0 min, 1% A; 11.0–14.0 min, 1% A; 14.0–14.5 min, 88% A; and 14.5–16.0 min, 88% A. Data acquisition occurred in both positive and negative ion modes, and the resulting data were processed with MetaboScape 4.0 software. Metabolites were identified by matching the detected compounds to entries in the MassBank of North America (MoNA) and LipidBlast databases.

### 2.4. DPPH• Scavenging Assay

The DPPH• scavenging activity was assessed using microplates at 490 nm. In brief, 100 µL of each ingredient was combined with 100 µL of a 0.2 mM DPPH• solution in methanol. The mixtures were incubated at room temperature for 30 minutes in the dark to prevent light-induced reactions. Absorbance was then measured at 490 nm using a Varioskan TM LUX microplate reader (Thermo-Fisher Scientific, Massachusetts, USA).

The DPPH• scavenging activity was calculated using the following formula (eq. 1):

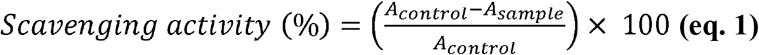

A_control_ represents the absorbance of the methanolic DPPH• solution without the sample and A_sample_ is the absorbance of the mixture containing the ingredient and DPPH• solution.

To determine the IC_50_ value, the scavenging activity (%) was plotted against the logarithmic concentration. A non-linear regression model with a five-parameter transformation was applied to estimate the concentration needed for 50% scavenging activity (Jaradat et al., 2021).

### 2.5. Oxygen Radical Absorbance Capacity (ORAC) Assay

The ORAC assay was performed using a spectrophotometric assay, utilizing AAPH as a peroxyl radical generator and Pyrogallol Red (PGR) as the colorimetric probe. Briefly, 194 µL of 75 mM phosphate buffer (pH 7.4), 15 µL of each ingredient, and 20 µL of 64 µM PGR were incubated at 37°C for 30 minutes. After incubation, 21 µL of 120 mM AAPH was added to each well, and absorbance was measured at 540 nm every 30 seconds over 3 hours.

The ORAC values were calculated using the following formula (eq. 2):

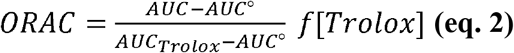

where AUC represents the area under the curve for PGR consumption in the presence of the ingredient, AUCº is the area under the curve without the ingredient, and AUC_Trolox_ is the area under the curve with Trolox. The dilution factor is denoted as *f*, and [Trolox] is the Trolox concentration in mM (Gregório et al., 2020).

### 2.6. Myoglobin Oxidation Assay

A 2% solution of equine heart myoglobin was prepared using a 50 mM phosphate buffer at pH 8.0. This myoglobin solution was then mixed with green tea extract or acerola juice powder at various concentrations (ranging from 50 to 1000 mg/L) by adding 20 µL of the antioxidant solutions to 180 µL of the myoglobin solution in a 96-well plate.

The absorbance of the myoglobin solution, both with and without antioxidants, was measured immediately at 503 nm, 525 nm, 557 nm, and 582 nm. After this initial measurement, the plates were incubated in the dark at three different temperatures: 25 °C and 35 °C. Absorbance readings at each wavelength were recorded every hour over a 24-hour period.

The oxidation states of myoglobin in the presence and absence of antioxidants were calculated using the method described by Jun et al. (2022), using the following equations (eqs. 4-11):

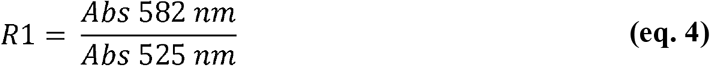

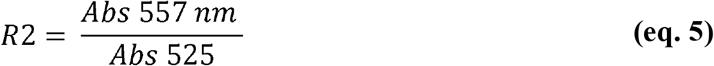

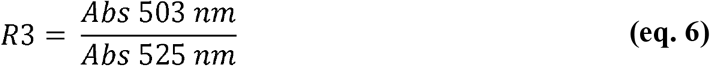

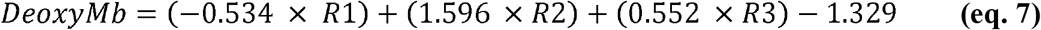

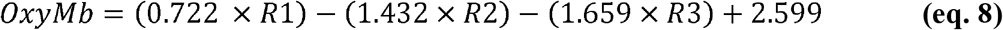

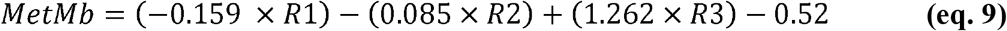

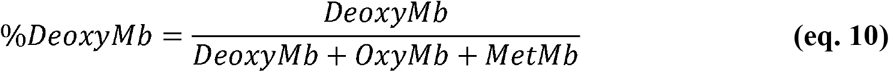

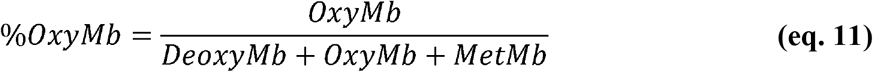

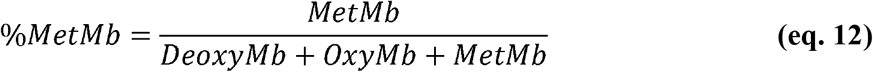

Where R1, R2 and R3 are absorbance value correlations for the different myoglobin forms: oxymyoglobin (OxyMb), deoxymyoglobin (DeoxyMb) and metmyoglobin (MetMb) respectively (Pujol et al., 2023).

Kinetic and thermodynamic parameters were determined with the following equations:

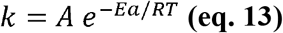

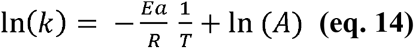

k is velocity constant, A the preexponential factor of Arrhenius, R gas constant and T temperature in K.

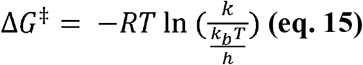

k_b_ is the Boltzmann constant, and h Planck constant

### 2.7. Statistical Analysis

The GraphPad Prism software, version 10.4.0, performed curve fitting and determined the area under the curve (AUC) for the collected data. A two-way ANOVA with Tukey’s multiple comparisons test was applied for the statistical analysis, with a significance level of *p* < 0.05.

## 3. Results

### 3.1. Antioxidant activity

The antioxidant activity of various food-grade botanicals was analyzed through DPPH radical scavenging and ORAC assays, including green tea, açai, acerola, beetroot, rosemary, and a blend of rosemary and green tea. The results showed significant differences in IC□□ values for DPPH radical scavenging capacity across the botanicals (Figure 1a). Beetroot exhibited the highest IC□□ value, around 2.0 µg/µmol DPPH, which was significantly higher than the other botanicals (p < 0.05), followed by açai with a value close to 1.5 µg/µmol DPPH. In contrast, green tea, rosemary, the rosemary, and green tea blend had low IC□□ values, all below 0.5 µg/µmol DPPH, with no significant differences (*p* > 0.05). Green tea showed an IC□□ value close to zero, highlighting its high antioxidant capacity in DPPH radical reduction (Figure 1a).

**Figure 1.**
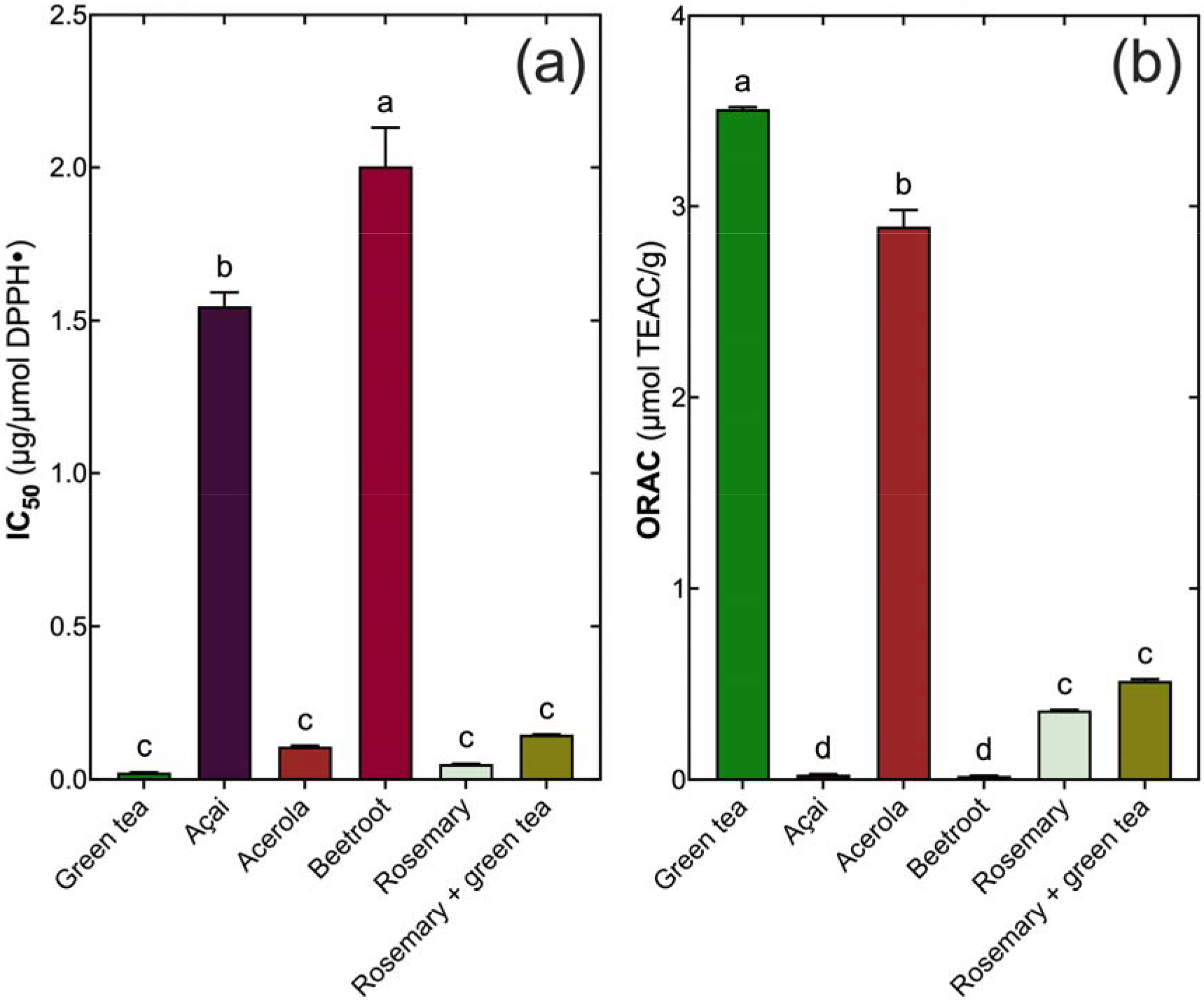
Antioxidant activity of food-grade botanicals. (a) DPPH radical scavenging capacity (IC□□ values in µg/µmol DPPH) and (b) ORAC (µmol TEAC/g). Bars represent mean ± standard deviation. Different letters above bars indicate significant differences between samples (*p* < 0.05).

For antioxidant activity measured through the ORAC assay, the results once again highlighted green tea as having the highest capacity, with an ORAC value of approximately 3.5 µmol TEAC/g, the highest among the botanicals tested (*p* < 0.05) (Figure 1b). Acerola followed with the second-highest ORAC value at about 3 µmol TEAC/g, and beetroot came next with around 2 µmol TEAC/g. Rosemary and the rosemary-green tea blend displayed intermediate ORAC values, close to 1 µmol TEAC/g with no significant differences among these three (*p* > 0.05) (Figure 1b). Açai, in contrast, had the lowest ORAC value, nearly zero, indicating minimal antioxidant activity in this assay compared to more effective botanicals like green tea and acerola (Figure 1b).

These findings reveal variations in the antioxidant capacity of the food-grade botanicals tested. Green tea and acerola had high activity levels in both assays, while açai demonstrated lower activity in the ORAC assay. Additionally, the higher IC□□ values for beetroot and açai in the DPPH assay suggest lower effectiveness in scavenging free radicals than other botanicals (Figure 1a and 1b).

### 3.2. Metabolite profiling

The metabolomic analysis of different food-grade botanicals revealed a diverse range of bioactive compounds, with concentrations shown in Figure 2. Each compound was quantified in green tea, açai, acerola, beetroot, rosemary, and a blend of rosemary and green tea, with values presented on a logarithmic concentration scale to highlight relative abundances across sources (Figure 2). The results showed significant variability in the concentration of specific metabolites among the different sources (Figure 2). Green tea stood out with high levels of phenolic compounds such as epicatechin, gallocatechin, epicatechin gallate, epigallocatechin, and epigallocatechin gallate, all of which were notably abundant compared to other sources (Figure 2). These compounds are well-known for their antioxidant activity, and their high concentrations underline green tea’s unique bioactive profile (Figure 2).

**Figure 2.**
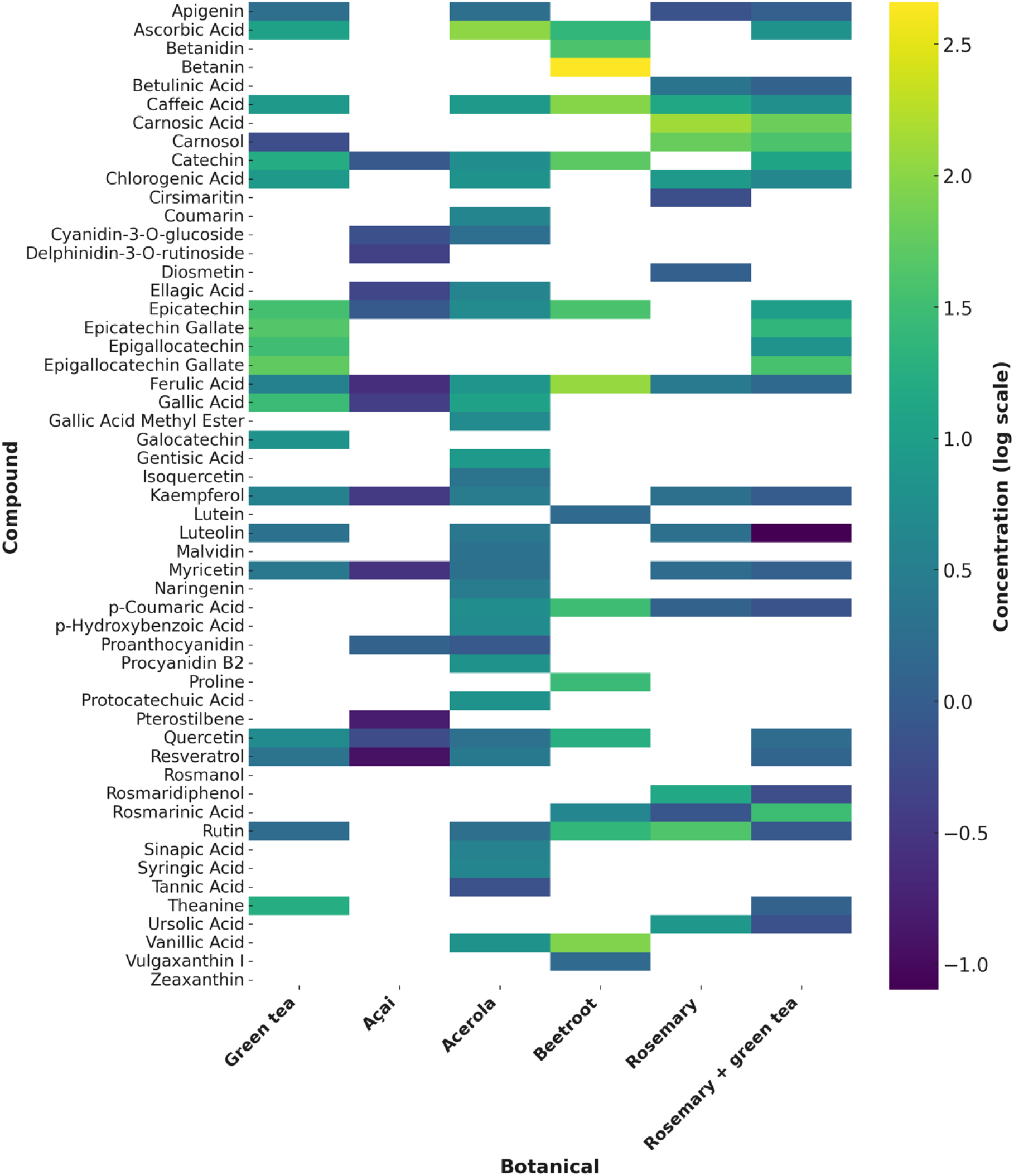
Concentration of bioactive compounds across different food-grade botanicals. Results include green tea, açai, acerola, beetroot, rosemary, and a blend of rosemary and green tea. The concentration of each compound is represented on a logarithmic scale to illustrate relative abundances.

In the case of açai, the most abundant compounds included ellagic acid, quercetin, and kaempferol (Figure 2). This profile suggests a composition rich in flavonoids and phenolic acids, which may also contribute to açai’s antioxidant capacity (Figure 2). Acerola, on the other hand, was distinguished by a high concentration of ascorbic acid, which was one of the most prevalent compounds in this source (Figure 2). In addition to ascorbic acid, acerola contained moderate levels of ferulic acid and myricetin, although in lower amounts than compounds specific to other sources (Figure 2).

Beetroot displayed a unique profile dominated by high concentrations of betanin and caffeic acid (Figure 2). Betanin, in particular, is a characteristic compound of beetroot and was present in much higher quantities than other compounds in this source (Figure 2). Additionally, moderate levels of p-coumaric acid were detected in beetroot, contrasting with its low or undetectable presence in the other sources (Figure 2). For rosemary, the dominant compounds included carnosic acid, carnosol, rosmarinic acid, and rosmanol (Figure 2). These phenolic diterpenes and related acids are characteristic of rosemary and were not observed in comparable concentrations in other sources (Figure 2). The rosemary and green tea blend also showed high levels of carnosic acid and carnosol, though in lower amounts than in pure rosemary, suggesting a possible dilution of these compounds in the blend (Figure 2).

On the other hand, gallic acid and rutin were detected across multiple sources, though in varying concentrations (Figure 2). Gallic acid was notably present in green tea, acerola, and rosemary, while rutin had a broader distribution, with significant levels in açai and beetroot (Figure 2). Other compounds, such as gentisic acid, luteolin, and naringenin, were found in smaller amounts and showed a more heterogeneous distribution across the different botanicals analyzed (Figure 2).

Regarding carotenoids and other pigments, zeaxanthin was detected in açai and lutein in rosemary, though both were present at relatively low concentrations compared to other compounds (Figure 2). Additionally, secondary metabolites like myricetin and resveratrol were identified in specific sources; myricetin was more abundant in acerola, while resveratrol was found at higher levels in the rosemary and green tea blend (Figure 2).

The distribution of compounds like protocatechuic acid, sinapic acid, and syringic acid also varied notably among the sources (Figure 2). Protocatechuic acid was relatively high in acerola, whereas sinapic acid and syringic acid were found in smaller quantities and with less consistency across the different sources (Figure 2). Trace amounts of ursolic acid were also detected in green tea and rosemary, though in low concentrations, suggesting that this triterpene is present only in limited quantities in these sources compared to other compounds (Figure 2).

These findings highlight the diversity and specificity of metabolomic profiles across the different food-grade botanicals (Figure 2). The structures and their origins are summarized in Figure 3.

**Figure 3.**
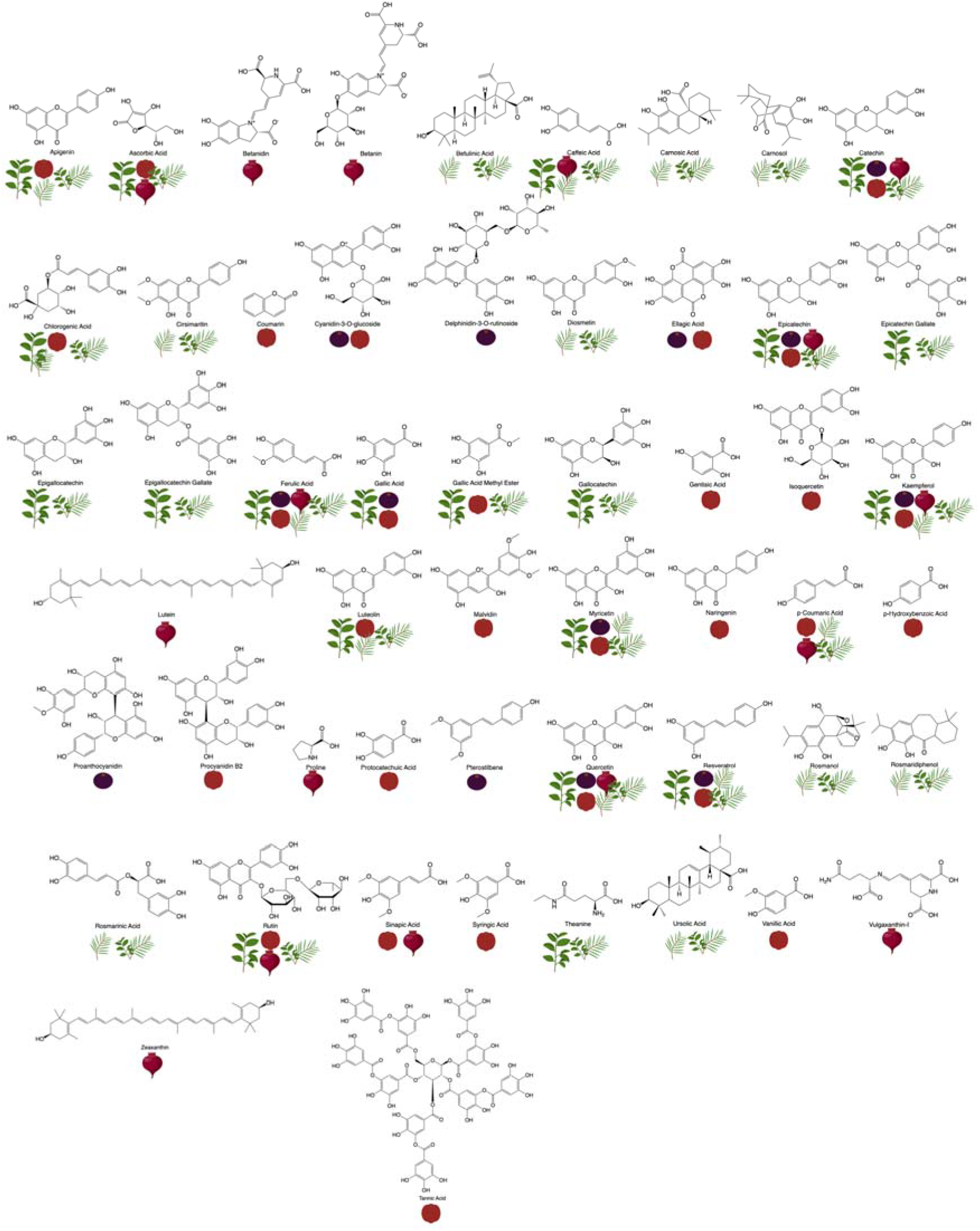
Chemical structures of bioactive compounds identified in food-grade botanicals. Botanicals include green tea, açai, acerola, beetroot, rosemary, and a rosemary-green tea blend. Icons below each structure indicate the botanical sources in which each compound was detected.

### 3.3. Myoglobin Oxidation Inhibition Kinetics

As shown in Figure 4, the inhibition rate of myoglobin oxidation (% inhibition/h/mg myoglobin/µg antioxidant) increased with the concentration of botanical extracts from green tea and acerola, tested at two temperatures (25 °C and 35 °C). These extracts were chosen for this study due to their impressive antioxidant performance in prior DPPH and ORAC assays, where both green tea and acerola demonstrated substantial free radical scavenging and oxygen radical absorbance capacities. This initial screening highlighted their potential as effective antioxidants, making them strong candidates for further analysis in this model.

**Figure 4.**
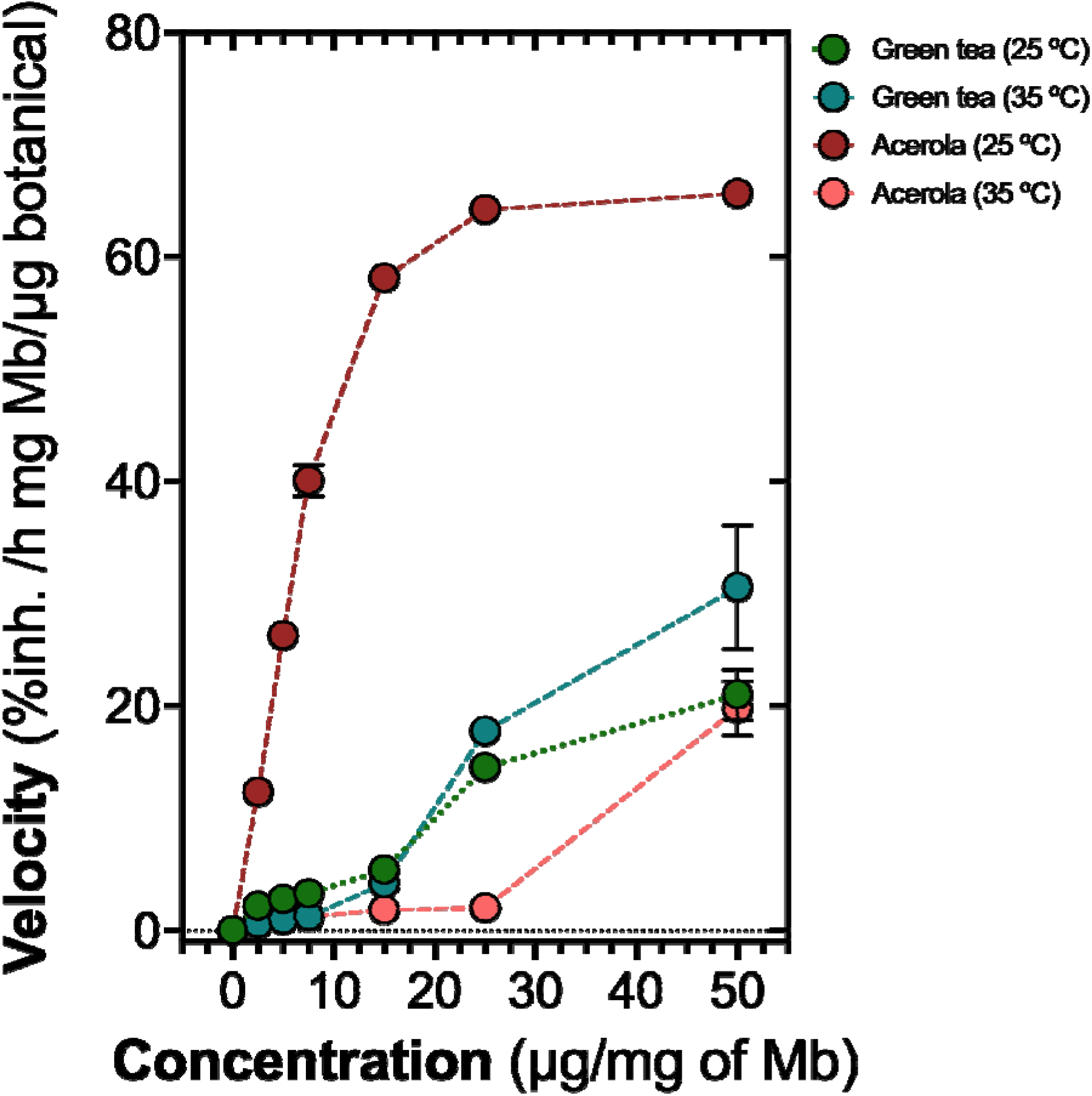
Inhibition rate of myoglobin oxidation (% inhibition/h/mg myoglobin/µg antioxidant) as a function of antioxidant concentration (µg/mg myoglobin) for green tea and acerola extracts at 25 °C and 35 °C. The extracts were selected based on their antioxidant capacity as determined by DPPH and ORAC assays. Data points represent the mean values (n = 3), and error bars indicate the standard deviation.

At concentrations below 20 µg/mg myoglobin, green tea and acerola displayed relatively low inhibition rates at both temperatures, with green tea at 25 °C showing minimal activity. However, as the antioxidant concentration increased, a distinct pattern emerged. The acerola extract at 25 °C showed a rapid rise in inhibition rate, reaching close to 60% inhibition/h at concentrations between 10 and 15 µg/mg myoglobin and leveling off at higher concentrations. This trend suggests a high efficacy for acerola at this temperature, allowing it to quickly reach its peak inhibitory effect.

In contrast, green tea exhibited a more gradual, temperature-dependent response. At 35 °C, green tea achieved higher inhibition rates compared to its performance at 25 °C, especially at concentrations above 20 µg/mg myoglobin, where the inhibition rate rose to approximately 25% inhibition/h. This increase in activity at higher temperatures suggests that green tea’s effectiveness may be enhanced by warmth, a result that aligns with the thermosensitivity of its antioxidant compounds, as seen in earlier DPPH and ORAC results.

On the other hand, acerola at 35 °C showed a less pronounced inhibition curve than at 25 °C, peaking at around 20% inhibition/h at a concentration of 50 µg/mg myoglobin. This indicates that acerola’s inhibitory effect is more potent at moderate temperatures, with a decline in efficacy as the temperature increases. These observations reflect acerola’s specific antioxidant profile from the DPPH and ORAC assays, suggesting that certain compounds in acerola perform best at lower temperatures. These findings underscore the distinct behaviors of green tea and acerola extracts, influenced by both concentration and temperature and highlight the importance of optimizing conditions to maximize antioxidant activity.

The kinetic and thermodynamic parameters for green tea and acerola extracts were calculated, showing distinct values for their activation energies and free energies of activation at different temperatures (Table I). The Ea for green tea was positive, with a value of 64.41 ± 9.47 kJ/mol, while the acerola extract displayed a negative Ea of -166.52 ± 5.75 kJ/mol, indicating a difference in the energy involved in the inhibition process for each extract.

**Table I.** Kinetic and thermodynamic parameters of activation energy (Ea) and free energy of activation (ΔG^‡^) for green tea and acerola extracts at 25 °C and 35 °C. Values are presented as mean ± standard deviation.

| Botanical | $E_a$ (kJ/mol) | $\Delta G^\ddagger$ (kJ/mol)<br>@ 25 °C | $\Delta G^\ddagger$ (kJ/mol)<br>@ 35 °C |
| --- | --- | --- | --- |
| Green tea | $64.41 \pm 9.47a$ | $-77.52 \pm 1.09a$ | $-82.37 \pm 3.85b$ |
| Acerola | $-166.52 \pm 5.75b$ | $-81.98 \pm 5.50b$ | $-79.23 \pm 5.92a$ |

Additionally, the ΔG^‡^ values were determined at 25 °C and 35 °C for both extracts. At 25 °C, green tea had a Delta G of -77.52 ± 1.09 kJ/mol, while the acerola extract reached - 81.98 ± 5.50 kJ/mol. At 35 °C, the ΔG^‡^ for green tea decreased to -82.37 ± 3.85 kJ/mol, whereas for acerola, it was -79.23 ± 5.92 kJ/mol.

These results indicate variation in each extract’s behavior with temperature, as seen in the calculated Ea and ΔG^‡^ values, highlighting an influence of temperature on the oxidation inhibition dynamics between green tea and acerola (Table I).

## Discussion

The study investigated the antioxidant capabilities of food-grade botanical extracts, specifically green tea, and acerola, in preventing the oxidation of myoglobin under varying temperature conditions. This research was driven by the growing need for natural antioxidant solutions in the food industry, especially considering consumer concerns over synthetic antioxidants and their potential health risks. The demand for natural alternatives is especially pertinent in plant-based products, where myoglobin or myoglobin-like substances enhance the appearance and appeal of meat analogs (Ahmad et al., 2022; Najmi et al., 2022).

The antioxidant efficacy of green tea and acerola extracts was first characterized through DPPH and ORAC assays, demonstrating substantial free radical scavenging and oxygen radical absorbance capacities for both extracts. Green tea exhibited the highest DPPH scavenging ability and the most elevated ORAC value among the botanicals tested. It suggests it is a robust antioxidant source rich in polyphenolic compounds like catechins known for their thermosensitive properties (Zhao et al., 2022). Similarly, acerola showed high antioxidant activity, with significant levels of ascorbic acid contributing to its radical scavenging potential (de Aquino Souza Miskinis et al., 2023). These initial findings underscored the viability of green tea and acerola as natural sources of antioxidants, warranting further examination of their effects on myoglobin oxidation.

The subsequent myoglobin oxidation assays revealed differential inhibition rates by the two extracts, which varied with concentration and temperature (de Aquino Souza Miskinis et al., 2023). At 25 °C, acerola showed a rapid rise in the inhibition rate, reaching nearly 60% inhibition per hour at low concentrations. This plateaued at higher concentrations, suggesting that acerola could quickly exert its maximum inhibitory effect at moderate temperatures (Kondjoyan et al., 2022). The efficacy of acerola at 25 °C aligns with its high ascorbic acid content, as vitamin C is an efficient, non-enzymatic antioxidant that stabilizes redox-sensitive proteins like myoglobin under controlled, moderate temperatures (Chung et al., 2024). This rapid saturation effect might reflect acerola’s ability to neutralize reactive species effectively at lower energy input levels, making it suitable for applications requiring stability at ambient conditions (de Aquino Souza Miskinis et al., 2023).

In contrast, green tea exhibited a more gradual inhibition profile, strongly influenced by temperature (Luo et al., 2020). At 35 °C, green tea achieved higher inhibition rates than its performance at 25 °C, with a noticeable increase in effectiveness at concentrations above 20 µg/mg of myoglobin. The temperature dependence of green tea’s antioxidant activity corresponds to the thermosensitivity of its primary polyphenols, including epigallocatechin gallate (EGCG) and other catechins (Sahu et al., 2024). These compounds are known to become more active with moderate heating, which can enhance their radical scavenging capabilities (Luo et al., 2020). However, high temperatures could eventually degrade these compounds, indicating that green tea’s inhibitory effects might be optimized in processes that involve mild heating but not extreme thermal exposure. The thermosensitivity observed suggests that green tea extracts could be strategically used in applications where slight temperature increases are expected, as it may enhance antioxidant potency without significant degradation.

When examining the kinetic and thermodynamic parameters, distinct behaviors were observed between green tea and acerola. The Ea for green tea was positive at 64.41 ± 9.47 kJ/mol, implying that higher temperatures facilitated its inhibitory effect on myoglobin oxidation (Duong et al., 2024; Pujol et al., 2023). This behavior is typical for compounds that benefit from moderate thermal activation, enabling more efficient radical scavenging and electron donation, especially in compounds with hydroxyl groups prone to oxidation at elevated temperatures (Ahmadi et al., 2020; Kondo et al., 1999). Conversely, acerola exhibited a negative Ea of -166.52 ± 5.75 kJ/mol, indicating a decrease in efficacy with temperature. This suggests that the antioxidant potential of acerola’s active compounds, such as ascorbic acid, may decrease under thermal stress, as vitamin C is known to degrade with heat exposure (Oey et al., 2006). This temperature sensitivity points to acerola’s suitability in applications where thermal stability is critical and can be incorporated without prolonged heating, preserving its bioactivity.

The ΔG^‡^ further illustrated the temperature-dependent dynamics of both extracts. For green tea, ΔG^‡^ values decreased from -77.52 ± 1.09 kJ/mol at 25 °C to -82.37 ± 3.85 kJ/mol at 35 °C, reflecting an increase in reaction spontaneity with temperature. This observation reinforces the hypothesis that green tea’s polyphenols become more reactive at slightly elevated temperatures, aligning with its performance in the oxidation inhibition assay. Meanwhile, acerola showed a decrease in ΔG^‡^ from -81.98 ± 5.50 kJ/mol at 25 °C to -79.23 ± 5.92 kJ/mol at 35 °C, indicating a slight reduction in inhibitory spontaneity with temperature. This pattern aligns with ascorbic acid’s known thermal degradation properties, which could lead to diminished antioxidant activity at higher temperatures. Combining these kinetic and thermodynamic parameters suggests that while green tea is suitable for applications involving mild heat, acerola’s effectiveness may be optimized in lower temperature settings, or its stability may need enhancement through encapsulation or other preservation techniques (Amorati et al., 2011; Duong et al., 2024; Galleano et al., 2010; Oey et al., 2006; Sahu et al., 2024).

The metabolomic profiling also provides context for these findings. Green tea’s profile was dominated by catechins such as epicatechin, gallocatechin, and EGCG, well-documented antioxidants with a strong affinity for scavenging free radicals and inhibiting lipid peroxidation (Luo et al., 2020; Zhao et al., 2022). The structural configuration of these catechins enables them to donate hydrogen atoms effectively, particularly under thermal activation, which explains the enhanced inhibition rates observed at 35 °C (Luo et al., 2020). Acerola’s metabolomic profile, by contrast, was rich in ascorbic acid and moderate levels of phenolic compounds like ferulic acid and myricetin (de Aquino Souza Miskinis et al., 2023; Duong et al., 2024). The predominance of ascorbic acid likely accounts for acerola’s rapid inhibitory effect at 25 °C, as vitamin C is known for its immediate electron-donating ability (Cort, 1974; Oey et al., 2006). However, its instability at higher temperatures may limit its use in heat-intensive processes.

The findings of this research also underscore the complexity of antioxidant behavior, which can vary significantly based on chemical composition and environmental conditions. Green tea’s gradual, temperature-responsive inhibition curve and high Ea value indicate that its polyphenols might be more suitable for formulations exposed to mild heating or temperature fluctuations, as this could enhance their antioxidant efficacy. In contrast, acerola’s rapid inhibition rate at lower temperatures and a negative Ea suggest it would be more effective in formulations designed for ambient or refrigerated storage, where its antioxidants remain stable. This differentiation is crucial for the food industry, where the selection of antioxidants often depends on each product’s specific storage and processing conditions (Colombo et al., 2020; Schilter et al., 2003).

In the context of myoglobin stabilization, these results have practical implications. For plant-based meat analogs or other applications where myoglobin or similar pigments are used to enhance visual appeal, selecting the appropriate antioxidant source can help maintain product quality over time (Devaere et al., 2022). For example, products intended for chilled storage might benefit more from acerola-based antioxidants (Duong et al., 2024). At the same time, those expected to undergo mild heating, such as during cooking or processing, might achieve better stability with green tea (Sahu et al., 2024). Furthermore, combining these extracts with other stabilizers could be explored to achieve a broader range of protection, as certain compound synergies might further enhance antioxidant efficacy across different temperature ranges.

Food-grade botanical antioxidants align with current consumer trends favoring natural, sustainable ingredients (Cao & Miao, 2023). As shown by this study, green tea and acerola both exhibit high antioxidant activity, and their efficacy in myoglobin stabilization suggests they could serve as viable replacements for synthetic antioxidants (Shang et al., 2020). Moreover, their diverse metabolomic profiles offer additional health benefits, as these compounds contribute to oxidative stability and the nutritional profile of food products (Poljsak et al., 2021).

In summary, this study demonstrated that green tea and acerola extracts exhibit distinct antioxidant behaviors based on temperature, concentration, and compound composition, influencing their effectiveness in myoglobin stabilization (Pujol et al., 2023). The data gathered provides valuable insights into optimizing antioxidant use in food applications, particularly for plant-based and natural products. These findings may guide future research and development of natural antioxidant formulations tailored to specific processing and storage conditions, offering the food industry natural alternatives that meet both functional and consumer-driven criteria.

## Conclusions

This study highlights the unique antioxidant properties of green tea and acerola extracts in preserving myoglobin stability. Acerola, rich in ascorbic acid, proved effective at moderate temperatures, showing rapid inhibition even at low concentrations. On the other hand, green tea displayed a temperature-sensitive inhibition pattern, with its activity increasing at 35 °C due to the heat sensitivity of its polyphenolic compounds. Further analysis of Ea and ΔG^**‡**^ confirmed these patterns: green tea’s positive Ea indicates it performs better with heat, while acerola’s negative Ea suggests reduced effectiveness at higher temperatures. These findings suggest green tea could be ideal for mildly heated food applications. At the same time, acerola may be better suited for ambient or cold storage, serving as a valuable natural alternative to synthetic antioxidants for color stability in food products.

## Conflict of Interest Statement

The authors confirm that no conflicts of interest could have influenced the development or results of this study.

## Notes

### Competing Interest Statement

The authors have declared no competing interest.

